# Uncovering kinesin dynamics in neurites with MINFLUX

**DOI:** 10.1101/2023.12.15.571866

**Authors:** Jan Otto Wirth, Eva-Maria Schentarra, Lukas Scheiderer, Victor Macarrón-Palacios, Miroslaw Tarnawski, Stefan W. Hell

**Author notes:** Authors contributed equally (Jan Otto Wirth, Eva-Maria Schentarra).

## Abstract

Neurons grow neurites of several tens of micrometers in length, necessitating active transport from the cell body by motor proteins. By tracking fluorophores as minimally invasive labels, MINFLUX is able to quantify the motion of those proteins with nanometer/millisecond resolution. Here we study the substeps of a truncated kinesin-1 mutant in primary rat hippocampal neurons, which have so far been mainly observed on microtubules polymerized on glass coverslips. A gentle fixation protocol largely maintains the structure and surface modifications of the microtubules in the cell. By analyzing the time between the substeps, we identify the ATP-binding state of kinesin-1 and observe the associated rotation of the kinesin-1 head in neurites. We also observed kinesin-1 switching microtubules mid-walk, highlighting the potential of MINFLUX to study the details of active cellular transport.

---

Kinesin-1 (herein referred to as kinesin) is a homodimeric motor protein of the kinesin superfamily known for taking distinct steps along single microtubule protofilaments. In a step, the motor domains (heads) move by 16 nm to the next free binding site. Kinesin has been studied extensively using fluorescence and scattering microscopy techniques ^1-6^, the latter having revealed that each step of the head consists of two ∼8 nm-sized substeps ^6-9^. While those scattering-based techniques exploit the high signal available from a laser, they come at the cost of requiring labels that are by an order of magnitude larger than the head itself. This size discrepancy raises the question whether the substeps are altered, or caused, by steric hindrance or other factors ^10,11^.

Interestingly, substeps of the head have recently been visualized using MINFLUX ^12,13^, a method that requires substantially fewer detected fluorescence photons for localizing single fluorophores than its established camera-based counterparts. MINFLUX localizes by probing with an excitation laser beam featuring a minimal (zero) intensity point or line at the center. Iterative matching of this minimal intensity center with the position of the fluorophore substantially reduces the number of photon detections required for attaining a certain localization precision, typically by about 100-fold ^14^. As a result, tracking single fluorophores with a spatio-temporal resolution of the order of nanometer/millisecond is readily attained. Harnessing the minimally impeding approach provided by direct fluorophore labeling, MINFLUX has recently been applied to study the conformational changes of kinesin walking on microtubules. An essential part of the cytoskeleton, microtubules are relatively stiff hollow polymers serving as tracks for motor proteins in the cell. It has been found that kinesin binds ATP between substeps while only one of the heads is bound to the polymer, i.e., during the one-head-bound (1HB) state, and that the heads undergo orientational changes during the substep.

However, all studies showing substeps – no matter if performed with small or large labels – have been conducted with microtubules polymerized on glass coverslips. Therefore, it remains uncertain if substeps do also occur on microtubules that were naturally assembled in cells. Although a recent MINFLUX study of kinesin in cells observed substeps ^15^, the observation was so scarce that the question remained whether the observed steps are rare by nature or just rarely observed due to limited spatio-temporal resolution or impediment by the label.

Here we show that substeps occur frequently and regularly in fixed primary rat hippocampal neurons (rHPNs) at up to physiological ATP concentrations. Moreover, we find an inverse relationship between the walking speed and the duration of the unbound state, providing evidence that ATP is taken up while only one head is microtubule-bound. Finally, in line with observations on microtubules polymerized on a glass coverslip, we observe label position dependent asymmetries in substep sizes as well as sideward displacements of kinesin walking on microtubules in neurites.

Microtubules polymerized in cells substantially differ from those polymerized on a coverslip, as they are decorated with microtubule-associated proteins (MAPs) and other biomolecules. Besides, tubulin may have undergone post-translational modifications. All these factors can act as roadblocks, forcing a motor protein to circumvent ^16-18^ the obstacle on its path. The way in which the cellular microtubules also affect the stepping behavior of kinesin remains unclear. Here we use gently fixed rHPNs, maintaining the native structure of the microtubules, as previously described ^19^. Neurons are highly relevant in this context, as they feature long and highly branched neurites, such as axons and dendrites, which require active transport of cellular cargo. Using fixed neurons, we employed purified motor proteins labeled at the head by maleimide coupling and controlled the ATP concentration of the buffer.

Since substeps have been shown on microtubules polymerized on coverslips for ATP concentrations ranging from 10 μM to 1 mM ^13^, and virtually no substeps have been found in cells at 5 mM ^15^, we measured at 50 μM, 500 μM and 5 mM thus covering the complete range from low-ATP up to physiological concentrations. The MINFLUX tracking routine was triggered by single motors walking into a stationary confocal volume on a neurite (Fig. 1a, top). The tracking routine consisted of repeated localizations in the x- and y-directions using line-shaped minima oriented along the x- and y-axis (Fig. 1a, bottom).

**Fig. 1.**
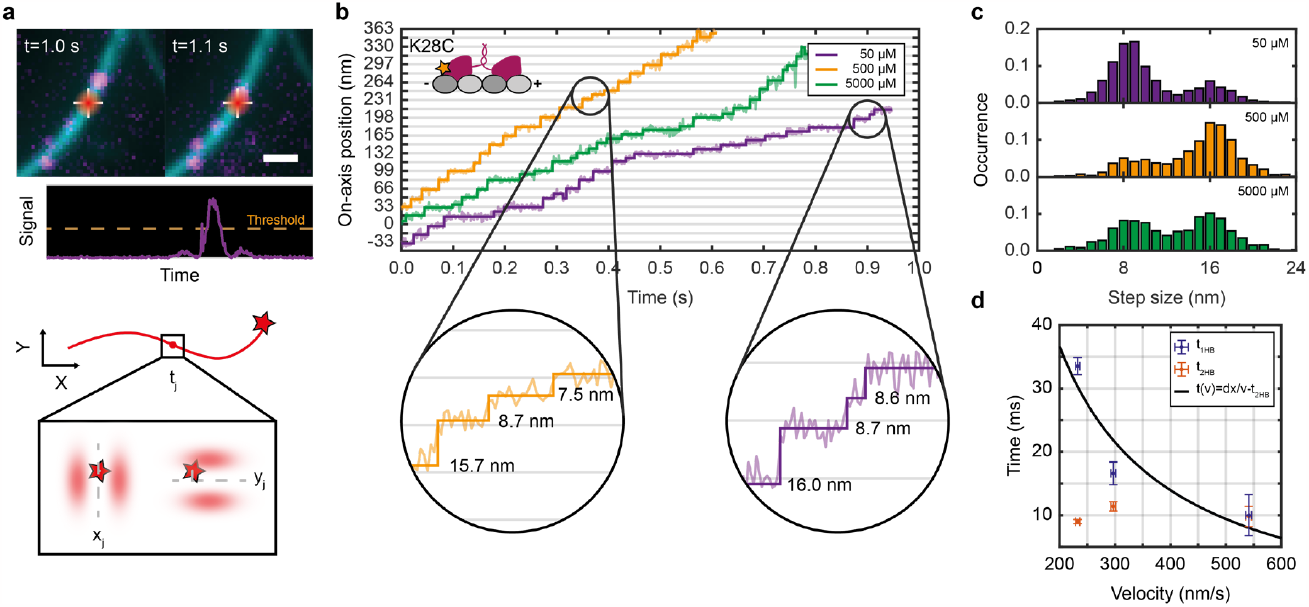
Visualization of substeps in primary rat hippocampal neurites. **(a)** Fluorophore-labeled kinesin motor walking into the stationary confocal volume creating an increased signal and triggering a MINFLUX tracking measurement (top). During the MINFLUX tracking process, for each time step t_j_ the x- and y-position of the minimum are updated to the newly estimated emitter position (bottom). **(b)** Exemplary on-axis position-time traces of construct K28C (inset) labelled at the back of the head domain recorded at 50 μM (purple), 500 μM (orange) and 5 mM (green) ATP concentration. The raw position data (semi-transparent lines) are overlaid with a step-fit (solid lines). Zoom-ins highlight ∼16 nm regular steps and pairs of similar-sized ∼8 nm substeps. **(c)** Population-normalized histograms of the measured step sizes recorded at the three ATP concentrations showing two populations of step sizes; 8 nm substeps and 16 nm regular steps (N_50μM_=1699, N_500μM_=1931, N_5mM_=1818). **(d)** Average duration of the one-head-bound (1HB) state between substeps (purple) and the two-head-bound (2HB) state (orange) plotted over the average velocity at 50 μM 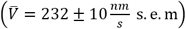,500 μM 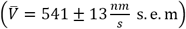 and 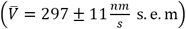. The solid black line shows a fit modeling the 1HB duration to velocity dependence by a simple rational function with parameters dx = 9.1 ± 5.8 nm and t_2HB_ = 8.8 ± 19.4 ms. Dwell times and velocity were averaged from in total 348 traces of 12 biological replicates. Error bars denote the standard deviation. Scale bar: 1 μm (a).

The first motor investigated was labeled at amino acid position 28 (construct K28C), which is located near the N-terminus at the back of the head with respect to the walking direction; see exemplary traces recorded at the three ATP concentrations (Fig. 1b). Traces were processed by an iterative step fit and substeps were identified by applying a Hidden-Markov-Model. This also allows discerning between the labeled head being microtubule-bound (bound state after 16 nm steps) or not (unbound state between paired substeps). The full ∼16 nm steps between binding sites are clearly visible and so are the substeps of ∼8 nm. The step size histograms disclose that the fraction of detected substeps depends on the ATP concentration, with most substeps being detected at 50 μM ATP and the least being detected at 500 μM (Fig. 1c). Relating the detected step sizes to the average walking speed of each trace reveals the fraction of substeps being velocity dependent, with higher walking speed resulting in fewer substeps detected (Supplementary Fig. S1). This observation is supported by evaluating the duration of the unbound state in-between paired substeps and the duration of the bound state in-between two full steps. From the histograms of these durations the average durations t_1HB_ and t_2HB_ that kinesin spends in a one-head-bound (1HB) state and a two-head-bound (2HB) state are extracted assuming equal kinetics of both heads (Supplementary Fig. S2). While no strong dependence on the walking speed can be observed for the 2HB state duration, the duration of the 1HB state monotonously decreases with increasing walking speed (Fig. 1d). Furthermore, since the walking speed is proportional to the ATP turnovers and thus the ATP binding rate, we find that ATP binding occurs during the 1HB state in the neurites as well.

Also, the walking speed does not saturate with increasing ATP levels, as usually described in the literature ^2,20,21^. We ascertained the decrease in walking speed at 5 mM ATP using fluorescence widefield microscopy, and even showed decreased kinesin activity at above 5 mM using an ATPase activity assay (Supplementary Fig. S3).

Next, we switched to a motor labeled at amino acid position 324 (construct T324C). The label is designed to be close to the neck linker. To compare the stepping behavior to that of construct K28C, we recorded traces at 50 uM and also at 5 mM ATP concentration in order to maximize the number of detected substeps (Fig. 2a). Both 16 nm-sized full steps as well as many substeps could clearly be resolved. However, the histogram of detected step sizes does now show a clear separation between full steps and substeps due to a rather broad substep distribution (Fig. 2b). To achieve an unbiased interpretation of this observation we plotted the size of consecutive steps against each other in a bivariate density scatter plot (Fig. 2c middle). The data clusters into four different regions centered around (16 nm, 16 nm), (16 nm, 8 nm), (8 nm, 8 nm) and (8 nm, 16 nm) with the greatest density at (16 nm. 16 nm) and (8 nm, 8 nm). As mentioned previously, we assume the labeled head to be microtubule-bound after a full step and unbound in-between paired substeps. Therefore substeps after a full step (16 nm, 8 nm) describe transitions from the bound to the unbound state which are paired with a substep before a full step (8 nm, 16 nm). Comparing the distributions of both constructs shows that, while the pairs of substeps of construct K28C uniformly spread around 8 nm, those of construct T324C are found to be in a range of 6 – 10 nm (combining to 16 nm) with the first substep being usually smaller than the second (Fig. 2c right).

**Fig. 2.**
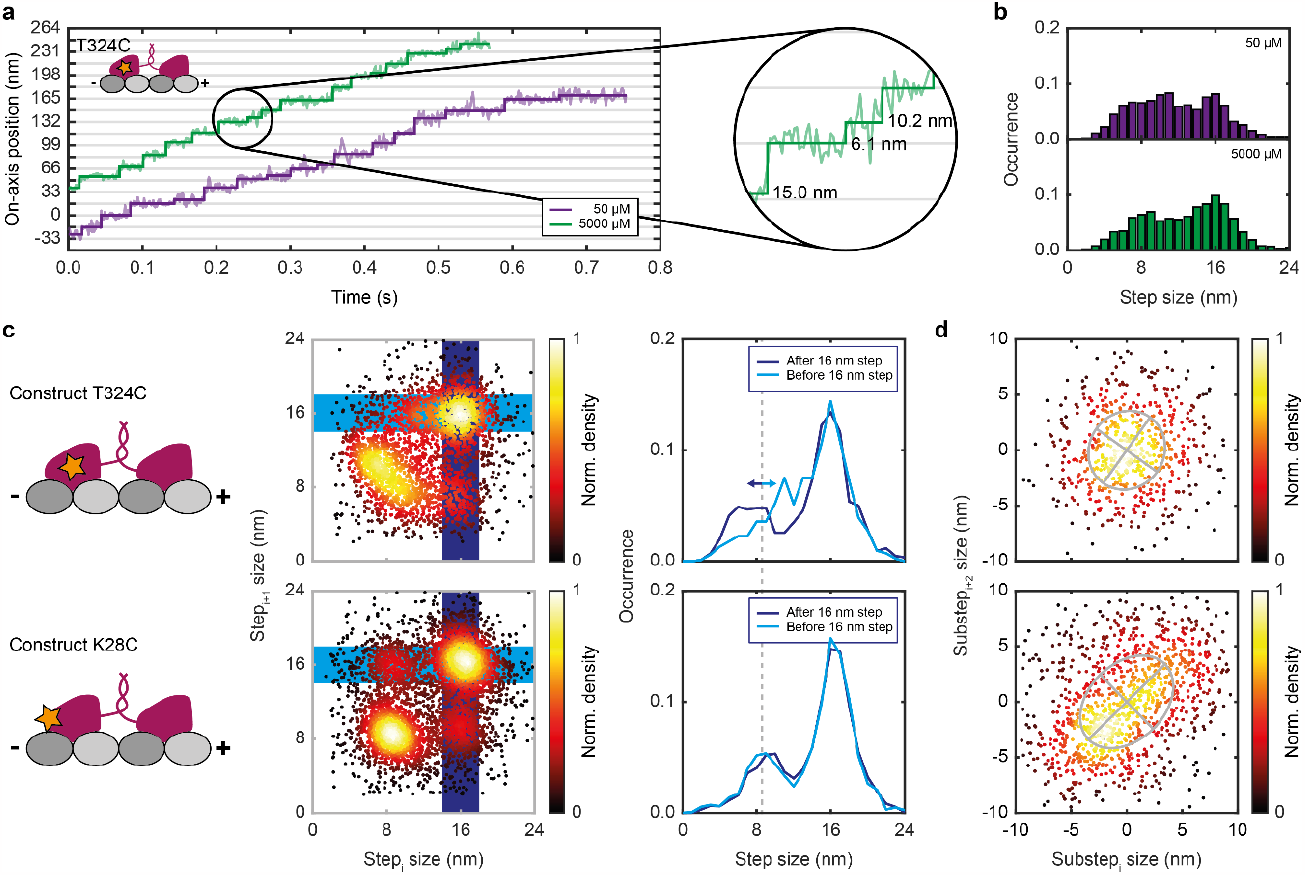
Comparison of the stepping behavior for constructs labeled at different amino acids. **(a)** Exemplary on-axis position-time traces of construct T324C (inset) labeled at the center-right of the head domain recorded at 50 μM (purple) and 5 mM (green) ATP concentration. The raw position data (semi-transparent lines) are overlaid with a step-fit (solid lines). The zoom-in highlights 16 nm regular steps and unequally-sized substeps. **(b)** Population-normalized histograms of measured step sizes recorded at the two ATP concentrations showing step sizes of 16 nm and substeps between 6 nm and 11 nm (N_50μM_*=*1625, N_5mM_=1592). **(c)** Comparison of the on-axis stepping behavior between construct T324C and K28C (constructs depicted on the left). The sequence of step sizes is displayed by a bivariate density scatter plot (middle). The step sizes recorded after a regular step are underlaid in dark blue, those before a regular step in light blue. The projections along the short axis of these boxes are shown as plots (right) showing similar substep sizes for construct K28C and varying substep sizes for construct T324C. The total number of steps for construct T324C and K28C are given by N_T324C_=3301 and N_K28C_=5519, respectively. **(d)** Comparison of the off-axis stepping behavior. Bivariate density scatter plots of the off-axis displacement for all subsequent bound to unbound and unbound to bound substeps together with the corresponding ellipses from the eigenvalues and eigenvectors of the covariance matrices (N_T324C_*=*590, N_K28C_*=*1055). Data for construct T324C is taken from 216 traces of 12 biological replicates (for statistics on construct K28C refer to Fig. 1).

We evaluated the sidewards displacement during the unbound state by plotting the off-axis step size of consecutive bound-to-unbound and unbound-to-bound transitions against each other in a bivariate density scatter plot (Fig. 2d). Here we reach the limit of our localization precision (4-5 nm standard deviation for single localizations) as the sidewards displacement is typically on the same scale or smaller. However, while no clear correlation can be observed for the off-axis step size of construct T324C, a small correlation can be observed for construct K28C, consistent with the previously observed rotation of the head during the unbound state ^13^.

Lastly, we observed kinesin switching microtubules mid-walk (Fig. 3a-d). Specifically, we saw kinesin motors displaying abrupt changes to their off-axis position and occasionally reversing their walking direction (Fig. 3b, c). As kinesin can only walk towards the plus-end of microtubules, no retrograde motors are present in the sample and the typical rate of recording traces was < 1 s^−1^, this phenomenon can most likely be explained by a microtubule switch. The observed switching occurred on small distances as well as between up to 80-nm-spaced microtubules, much larger than the typical 8 nm spacing between the heads. One of the examples (Fig. 3c), even exhibits two switches (at 0.3 s and 0.8 s) within a single trace. As the data represents a 2D-projection of the actual 3D movement, we assume the first switching was to a microtubule separated in the z-direction.

**Fig. 3.**
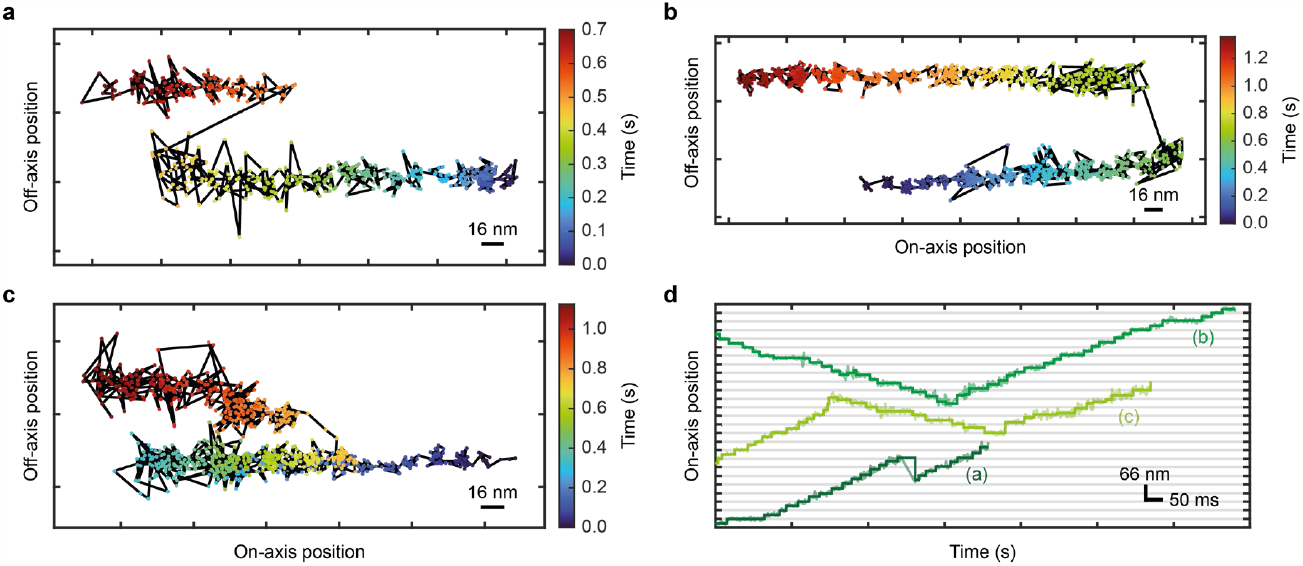
Kinesin switches microtubules and changes walking direction in the cellular context. **(a-c)** Exemplary 2D position traces of construct K28C color-coded for time, all displaying an off-axis displacement larger than the ∼25 nm diameter of a microtubule, indicating mid-walk microtubule switching. Individual localizations (points) are connected by black lines. The traces displayed in (a-b) show an off-axis displacement of ∼60 nm within 15-50 ms. While trace (a) continues progressing into the same direction, trace (b) switches to walking into roughly the opposite direction. The trace shown in (c) exhibits microtubule switching with ∼60 nm off-axis displacement within less than 5 ms, accompanied by a previous change in walking direction without off-axis displacement (indicated by the time color-coding). **(d)** Position-time traces of the on-axis movement of the traces shown in (a-c). The raw data (semi-transparent lines) are overlaid by the step fit (solid lines).

In conclusion, MINFLUX directly visualizes kinesin substeps on microtubules of lightly fixed neurites in a cellular context. While we validated the stepping characteristics that were recently reported for microtubules polymerized on a coverslip in neurites, the higher microtubule density in axons and dendrites allowed us to observe additional effects, such as microtubule switching. The recorded changes in the walking pattern of motors labeled at different positions highlight the importance of pursuing adequate and possibly multiple labeling strategies to minimize the risk of drawing conclusion influenced by the position and flexibility of the fluorophore. In any case, MINFLUX tracks the largely unimpeded motion of motor proteins in unprecedented detail, enabling the unraveling of protein dynamics and conformational changes directly in a cell.

## Acknowledgements

We thank Jasmine Hubrich, Jana Kress, Birgit Koch, and Alena Fischer for their work and support in the isolation and culturing of primary neurons, and particularly Jasmine Hubrich and Alena Fischer for their advice regarding the handling and labeling of neurons. We also thank Dr. Richard Lincoln for helpful feedback and critical reading.

## Author contributions

S. W. H. initiated the project. E.-M. S. prepared samples, performed measurements and evaluated data supervised by J. O. W. who also improved software for the MINFLUX tracking routine and the data evaluation. V. M.-P. supported and guided the culturing, fixation and labeling of neurons. L. S. assisted in validating motor activity and performed the activity assay together with E.-M. S.. M. T. did the cloning, expression and purification of the kinesin constructs. J. O. W. and E.-M. S. interpreted the results supported by critical discussions with all authors. J. O. W., E.-M. S. and S. W. H. wrote the manuscript with input from all authors.

## References

1 Svoboda, K., Schmidt, C. F., Schnapp, B. J. & Block, S. M. Direct observation of kinesin stepping by optical trapping interferometry. Nature 365, 721–727, doi:10.1038/365721a0 (1993).

2 Schnitzer, M. J. & Block, S. M. Kinesin hydrolyses one ATP per 8-nm step. Nature 388, 386–390, doi:10.1038/41111 (1997).

3 Hua, W., Chung, J. & Gelles, J. Distinguishing inchworm and hand-over-hand processive kinesin movement by neck rotation measurements. Science 295, 844–848, doi:DOI 10.1126/science.1063089 (2002).

4 Yildiz, A., Tomishige, M., Vale, R. D. & Selvin, P. R. Kinesin Walks Hand-Over-Hand. Science 303, 676–678, doi:doi:10.1126/science.1093753 (2004).

5 Mori, T., Vale, R. D. & Tomishige, M. How kinesin waits between steps. Nature 450, 750–U715, doi:10.1038/nature06346 (2007).

6 Verbrugge, S., Lansky, Z. & Peterman, E. J. Kinesin’s step dissected with single-motor FRET. Proc Natl Acad Sci U S A 106, 17741–17746, doi:10.1073/pnas.0905177106 (2009).

7 Mickolajczyk, K. J. et al. Kinetics of nucleotide-dependent structural transitions in the kinesin-1 hydrolysis cycle. Proc Natl Acad Sci U S A 112, E7186–7193, doi:10.1073/pnas.1517638112 (2015).

8 Isojima, H., Iino, R., Niitani, Y., Noji, H. & Tomishige, M. Direct observation of intermediate states during the stepping motion of kinesin-1. Nat Chem Biol 12, 290–+, doi:10.1038/Nchembio.2028 (2016).

9 Sudhakar, S. et al. Germanium nanospheres for ultraresolution picotensiometry of kinesin motors. Science 371, eabd9944. doi:doi:10.1126/science.abd9944 (2021).

10 Mickolajczyk, K. J., Cook, A. S. I., Jevtha, J. P., Fricks, J. & Hancock, W. O. Insights into Kinesin-1 Stepping from Simulations and Tracking of Gold Nanoparticle-Labeled Motors. Biophys J 117, 331–345, doi:10.1016/j.bpj.2019.06.010 (2019).

11 Hasnain, S., Mugnai, M. L. & Thirumalai, D. Effects of Gold Nanoparticles on the Stepping Trajectories of Kinesin. J Phys Chem B 125, 10432–10444, doi:10.1021/acs.jpcb.1c02218 (2021).

12 Balzarotti, F. et al. Nanometer resolution imaging and tracking of fluorescent molecules with minimal photon fluxes. Science 355, 606–612, doi:doi:10.1126/science.aak9913 (2017).

13 Wolff, J. O. et al. MINFLUX dissects the unimpeded walking of kinesin-1. Science 379, 1004–1010, doi:10.1126/science.ade2650 (2023).

14 Eilers, Y., Ta, H., Gwosch, K. C., Balzarotti, F. & Hell, S. W. MINFLUX monitors rapid molecular jumps with superior spatiotemporal resolution. Proceedings of the National Academy of Sciences 115, 6117–6122, doi:doi:10.1073/pnas.1801672115 (2018).

15 Deguchi, T. et al. Direct observation of motor protein stepping in living cells using MINFLUX. Science 379, 1010–1015, doi:10.1126/science.ade2676 (2023).

16 Schneider, R., Korten, T., Walter, W. J. & Diez, S. Kinesin-1 Motors Can Circumvent Permanent Roadblocks by Side-Shifting to Neighboring Protofilaments. Biophys J 108, 2249–2257, doi:10.1016/j.bpj.2015.03.048 (2015).

17 Dreblow, K., Kalchishkova, N. & Bohm, K. J. Kinesin passing permanent blockages along its protofilament track. Biochem Biophys Res Commun 395, 490–495, doi:10.1016/j.bbrc.2010.04.035 (2010).

18 Schmidt, C. et al. Tuning the “roadblock” effect in kinesin-based transport. Nano Lett 12, 3466–3471, doi:10.1021/nl300936j (2012).

19 Tas, R. P. et al. Differentiation between Oppositely Oriented Microtubules Controls Polarized Neuronal Transport. Neuron 96, 1264–1271 e1265, doi:10.1016/j.neuron.2017.11.018 (2017).

20 Verbrugge, S., van den Wildenberg, S. M. & Peterman, E. J. Novel ways to determine kinesin-1’s run length and randomness using fluorescence microscopy. Biophys J 97, 2287–2294, doi:10.1016/j.bpj.2009.08.001 (2009).

21 Visscher, K., Schnitzer, M. J. & Block, S. M. Single kinesin molecules studied with a molecular force clamp. Nature 400, 184–189, doi:10.1038/22146 (1999).

