## Supplements for "Uncovering kinesin dynamics in neurites with MINFLUX"

[†] Authors contributed equally

### 1. Materials and Methods

#### 1.1. Primary rat hippocampal neuron culture

Primary rat hippocampal neurons (rHPNs) were prepared from postnatal P0-P2 Wistar rats (Janvier Labs, Le Genest-Saint-Isle, France) of either sex with no genetic modification as previously described (*1*). Animals were sacrificed according to the guidelines established by the German Animal Welfare Act (Tierschutzgesetz der Bundesrepublik Deutschland, TierSchG) and the Animal Welfare Laboratory Animal Ordinate (Tierschutz-Versuchstierverordnung: TierSchVersV), following all regulations, conducted and documented under supervision of animal welfare officers of Max Planck Institute for Medical Research (MPImf) (permit number assigned by the MPImF: MPI/T-35/18). According to the TierSchG and the Tierschutzversuchstierverordnung no ethical approval from the ethics committee is required for the procedure of sacrificing rodents for subsequent extraction of tissues, as performed in this study. Briefly, hippocampi were collected and dissected all together from the same litter, digested with 0.25 % trypsin (Thermo Fisher Scientific, Waltham, MA, USA) for 20 min at 37 °C and subsequently dissociated and maintained in cell culture medium (Neurobasal™ supplemented with 2 % B-27, 1 % GlutaMAX and 1 % penicillin/streptomycin; all from Gibco, Thermo Fisher Scientific). Dissociated cells were plated at a density of 60,000 cells/well in 12-well plates on 18 mm coverslips pre-coated with 0.1 mg/ml poly-ornithine (Sigma-Aldrich/Merck, Darmstadt, Germany) and 1 µg/ml laminin (Corning, New York, NW, USA) in culture medium. Medium was exchanged after attachment of the cells (approximately 2 h) and neurons were maintained at 37 °C in a humidified incubator (95% rH) with 5 % $CO_2$ until 11 - 21 days in vitro (DIV).

#### 1.2. Constructs

Truncated human kinesin-1 (residues 1-560 with a C-terminal His-tag) with all solvent-exposed cysteines mutated to alanine (C7S, C65A, C168A, C174S, C294A, C330S, C421A) and containing a unique cysteine residue for labeling at position T324C was expressed using plasmid K560CLM T324C (*2*) (obtained from Addgene, #24460). A 'cysteine-light' truncated human kinesin-1 for labeling at K28C position (K560CLM K28C) was generated using QuikChange II (Agilent) site-directed mutagenesis method with CLM RP HTR plasmid (*3*) obtained from Addgene (#24430) and used as a template. The construct was verified by Sanger sequencing (Eurofins).

#### 1.3. Protein expression and purification

The vectors were transformed into E. coli BL21 CodonPlus(DE3)-RIL (Agilent). Cells were grown at 37 °C in LB medium supplemented with ampicillin (100 µg/ml) and chloramphenicol (30 µg/ml). At an optical density at 600 nm of 0.8-1.0, cells were transferred to 18 °C and expression was induced with 0.1 mM isopropyl β-D-1-thiogalactopyranoside (IPTG). Cells were harvested after overnight expression by centrifugation, frozen in liquid nitrogen and stored at -80 °C until further use. All subsequent purification steps were done at 4 °C.

The cell pellets were resuspended in lysis buffer (50 mM NaH2PO4 pH 8.0, 250 mM NaCl, 20 mM imidazole, 2 mM MgCl2) supplemented with cOmplete EDTA-free protease inhibitors (Roche), 20 µg/ml DNaseI, 10 mM β-mercaptoethanol, 1 mM ATP. The cells were lysed using a microfluidizer (Microfluidics) operated at a pressure of 0.9 MPa and the lysates were clarified by centrifugation at 47,850 g for 1 hour at 4 °C. The cleared supernatants were loaded onto a HisTrap FF 5 ml (Cytiva) column pre-equilibrated with lysis buffer. The column was washed with 50 mM NaH2PO4 pH 6.0, 250 mM NaCl, 20 mM imidazole, 1 mM MgCl2, 10 mM β-mercaptoethanol, 0.1 mM ATP and the protein was eluted with 50 mM NaH2PO4 pH 7.2, 250 mM NaCl, 500 mM imidazole, 1 mM MgCl2, 10 mM β-mercaptoethanol, 0.1 mM ATP.

Fractions containing kinesin were 5-fold diluted with buffer A (25 mM PIPES pH 6.8, 2 mM MgCl2, 1 mM EGTA, 0.2 mM TCEP, 0.1 mM ATP) before loading onto a HiTrapQ FF 5ml (Cytiva) column pre-equilibrated with buffer A containing 100 mM NaCl. The column was washed with the same buffer and the protein was subsequently eluted using linear gradient of 100-1000 mM NaCl in buffer A. The peak fractions containing kinesin were concentrated using Amicon Ultra (Merck Millipore) centrifugal units and further purified by gel filtration on a HiLoad 16/600 Superdex 200 pg (Cytiva) column equilibrated with buffer B (25 mM PIPES pH 6.8, 300 mM NaCl, 2 mM MgCl2, 1 mM EGTA, 0.2 mM TCEP, 0.1 mM ATP) to further improve the sample quality. Selected fractions containing kinesin were finally combined and concentrated, supplemented with 10 % (w/v) sucrose, aliquoted, frozen in liquid nitrogen and stored at -80 °C. Purified proteins were analyzed by ESI-MS.

##### 1.4. Labeling

Exposed solvent cysteines were labeled with ATTO 647N maleimide (AD 647N-41, ATTO-TEC) over night at 4 °C and excess dye was removed by size-exclusion chromatography (PD MiniTrap G-25, 28-9180-07, Cytiva). The degree of labeling was determined wit UV-Vis spectroscopy (DS-11+ Spectrophotometer, DeNovix) and mass spectrometry (ESI, maXis II ETD, Bruker) and aliquots of purified labeled kinesin were flash frozen in PEM80 (80 mM PIPES, 0.5 mM EGTA, 2 mM MgCl2 set to pH 7,40) supplemented with 10 % (w/v) sucrose and stored at −80 °C until use.

##### 1.5. Cell fixation

To prepare neurons for tracking, the cytoplasm was extracted by applying prewarmed 37 °C extraction buffer (1 M sucrose and 0.15 % (v/v) Triton-X-100 (AppliChem GmbH) in BRB80: 80 mM PIPES, 1 mM EGTA, 1 mM $MgCl_2$ set to pH 6.80) for 1 min. Next, the buffer was exchanged for prewarmed fixation buffer (0.5 M sucrose, 0.075 % (v/v) Triton-X-100 and 1 % (v/v) paraformaldehyde (PFA, Carl Roth) in BRB80) for 1 min. Subsequently the solution was exchanged for prewarmed washing buffer (1 µM paclitaxel (Focus Biomolecules) in BRB80) and incubated for 1 min. The washing step was repeated three times. Fixed microtubules were stained with BioTracker™ 488 Green Microtubule Cytoskeleton Dye (SCT142, Sigma-Aldrich/Merck; 1:1000 in BRB80) and incubated for at least 15 min at 37 °C. The staining solution was removed and the cells were blocked for 3 min with blocking buffer (2 mM ascorbic acid (BioVision Inc), 0.5 mg/ml casein (VWR Chemicals), 538 µg/ml catalase, 1 mM 1,4-dithiothreitol (DTT, Carl Roth), 41.7 µg/ml glucose oxidase, 1.7 % (w/v) glucose, 1 µM paclitaxel, 2 mM methyl viologen and 50 µM/500 µM/5 mM ATP; (all from Sigma-Aldrich/Merck unless stated otherwise) in BRB80). Finally coverslips with fixed cells were mounted on cavity slides (1320002, Paul Marienfeld GmbH & Co.KG, Lauda Königshofen, Germany) containing approximately 150 µL imaging buffer (2 mM ascorbic acid, 0.2 mg/ml casein, 538 µg/ml catalase, 1 mM DTT, 41.7 µg/ml glucose oxidase, 1.7 % (w/v) glucose, 1 µM paclitaxel, 2 mM methyl viologen and 50 µM/500 µM/5 mM ATP in BRB80) supplemented with 4-8 nM kinesin and edges were sealed with picodent twin-sil speed 22 (picodent Dental-Produktions- und Vertriebs GmbH) and cured for 5 min at 37 °C.

#### 1.6. MINFLUX tracking routine

Before recording MINFLUX position traces of kinesin, suitable microtubule regions were selected by performing 5×5 µm$^2$ confocal xy-scans of the 488 nm laser (pixel size 50 nm). Typically, thin and straight bundles were selected, with minimal background and devoid of overlapping processes. Up to 30 pixels were manually selected in a zig-zag arrangement over the width of a microtubule bundle and sequentially addressed by the galvo-scanner with 10 ms exposures of the 642 nm laser. Once the detected photon rate exceeded a predefined threshold ranging from 6 – 10 kHz, depending on the background of an individual sample, the MINFLUX tracking routine was triggered. During the initial zoom-in phase of the routine, a series of three MINFLUX steps with progressively decreasing L were performed, probing the excitation intensity minimum around the latest estimated molecule localization to positions $[-L/2, 0, L/2]$ in both dimensions. Following each cycle, the molecule's position estimate $x$ was updated based on the recorded photons according to

$$x_{new} = x_{old} + \frac{L}{4} \frac{n_- - n_+}{n_+ + n_- - 2n_0} \tag{1}$$

with $[n_-, n_0, n_+]$ being the photons recorded at $[-L/2, 0, L/2]$ respectively. Localizations are considered failed if either less than five photons are collected or if $2(n_+ + n_- - 2n_0) < |n_- - n_+|$ which limits the allowed corrections to $L/2$. Repeated localizations with the same $L$ ensure that no motor is missed the localization process due to a single failed localization. As the first three steps only ensure the centering of the dye, most repeats are performed in the last step. At this step $L$ was set to 50 nm for all measurements. For each dimension a localization consists of three exposures with an exposure time of 200 µs. Considering an additional dead time of 5 µs for switching between exposures and 500 ps for recalculating the updated molecule position, a single two-dimensional (2D) localization required 1231 µs. An overview over the $L$-values, laser powers and repeats is given in Table S1.

#### 1.7. TIRF/Widefield microscopy

Kinesin velocity measurements were performed on a custom widefield setup built around an Olympus IX83 microscope body (Olympus Corp., Tokyo, Japan) providing a motorized stage and a piezo Z-stage. The system is equipped with two excitation laser lines at wavelength 488 nm (Oxxius, Lannion, France) and 640 nm (Cobolt AB, Solda, Sweden). For image acquisition, the excitation laser was focused in the back focal plane of the 1.49 NA 100× oil objective lens (UAPON100XOTIRF, Olympus Corp.) controlled by an acousto optical tunable filter (AOTF, AA OPTO-ELECTRONIC, Orsay, France) and the emitted light was passed via a quad-band emission filter (F66-887, AHF analysentechnik AG, Tübingen, Germany) for separation of excitation and emission bands, respectively. Total internal reflection fluorescence (TIRF, Fish, 2009) illumination mode was achieved by moving the incident laser beam via the motorized stage to the outer part of the pupil plane. Emitted photons from the sample were captured by an iXon 897 EMCCD camera (Andor Technology, Belfast, UK). Active z-stabilization was provided based on the Olympus focus stabilization system (Confocus) and the microscope was operated by a custom-written LabView (National Instruments Corp., Austin, TX, USA) program.

Frame stacks were recorded at 50 µM, 500 µM and 5000 µM ATP concentration in 3 biological samples each. For each condition, 4 cells were imaged and time lapse acquisitions of 200 time points with an exposure time of 100 ms were captured. Moving kinesins (construct K28C) labeled with ATTO 647N on fixed neuronal microtubules were recorded with the 642 nm laser set to 5 mW laser power. (176 W/cm$^2$ focal intensity) Microtubules were imaged using the 480 nm laser at 1 mW laser power (353 W/cm$^2$ focal intensity). Frame series were analyzed in Fiji (*4*) determining the run length along microtubules for individual motors. Velocities were calculated from determined run-lengths over time.

#### 1.8. ATPase activity assay

Kinesin's ATPase activity of the construct K28C was determined at different ATP concentrations by using the commercial CytoPhos™ ATPase Endpoint assay (Cytoskeleton, Inc., DENVER, CO, USA) according to the manufacturer protocol. Briefly, 1 µg of kinesin was added to a solution of pre-stabilized microtubules (20 µM paclitaxel) with a final concentration of ~ 67 µg/ml, in imaging buffer without ATP. ATP was then added at concentrations of 10 µM, 50 µM, 100 µM, 500 µM, 1000 µM, 50000 µM and 10000 µM and after 15 min at room temperature, the reaction was stopped by adding the commercial CytoPhos™ reagent. The absorption at 650 nm was recorded in a plate reader after 10 min. To determine the kinesins-dependent amount of produced inorganic phosphate, CytoPhos™ reagent was added to known amounts of phosphate in MilliQ water an the absorption spectra at 650 nm were recorded and plotted as linear calibration curve over the employed amounts of phosphate.

#### 1.9. Data analysis

MINFLUX position traces of kinesin were screened and selected in LabView 2019 based on a sufficient number of detected photons for a successful localization, as well as them displaying a clear stepping behaviour and a good average SBR. Those traces were post-processed and analysed using dedicated Matlab scripts. First, the start and end of each trace are manually selected. Then, using a so-called sliding curvature estimator

$$x_{SCE}(t) = \frac{L}{4} \frac{n_-(t) - n_+(t)}{1/T \int_{t'-T/2}^{t'+T/2} n_+(t') + n_-(t') - 2n_0(t')dt'} \, , \qquad (2)$$

the position estimate was refined by averaging the curvature of the excitation pattern at each time t in a sliding time interval of around $T = 20$ ms. Typically, kinesin does not move within this time interval, thus validating to assume the curvature constant. By doing so, noise on the position estimate can be reduced, improving the localization precision without loss of temporal resolution. Next, the main movement of the trace is aligned with the x-axis. This so-called on-axis x-position is subjected to the step fit function described in (*5*), which is based on an iterative change point detection (*6*). The step function is additionally filtered by a moving median of width 9 (to remove steps caused by spikes in the data) and in a single round, steps below 2 nm are removed. Note that removing steps changes the plateau levels and thus affects the following or previous step and can cause them to become less than 2 nm. The change points of the on-axis step fit are then used to create the step fit for the off-axis position. From the step fits, the step sizes as well as the dwell time between steps are extracted.

In order to extract the dwell times between substeps and regular steps, as well as to correlate the off-axis displacement between substeps, the sequence of steps from single traces are analysed using a Hidden Markow model (*5*). The model identifies the most likely sequence of the labeled head being bound or unbound based on the assumptions that regular steps are 16 nm-sized and that substeps occur in pairs each being around 8 nm in size.

Additionally, the average localization precision and photon counts within each plateau are extracted as well as the run length and run time, from which the velocity can be calculated. All this information is saved in an excel sheet for pooled processing.

To further evaluate the average residence time kinesin spends in the one-head-bound (1HB) and two-head-bound (2HB) state the dwell times of the bound and unbound state pooled from all traces recorded with the same construct and ATP concentration are fitted simultaneously using the model derived in (*5*).

$$p_U(t) = \frac{e^{-\frac{t}{\tau_{1HB}}}}{\tau_{1HB}}$$

$$p_B(t) = \frac{1}{\tau_{2HB}(\tau_{2HB} - \tau_{1HB})}\left(\tau e^{-\frac{\tau}{\tau_{2HB}}} - \frac{\tau_{1HB}\tau_{2HB}}{\tau_{2HB} - \tau_{1HB}}\left(e^{-\frac{\tau}{\tau_{2HB}}} - e^{-\frac{\tau}{\tau_{1HB}}}\right)\right) \quad (3)$$

#### 1.10. Data representation

Traces illustrate the post-processed MINFLUX tracking position either over time together with their corresponding step function.

Histograms show the counts per individual bin normalized to the size of the population. Single datasets are displayed as bars and multiple datasets in a single plot as lines.

Scatter plots present either a sequence of step sizes, or xy-coordinates of a trace. For a sequence of step sizes each point is color-coded according to the number of points within a 4 nm radius. For xy-traces each point is color-coded for time.

Violin plots represent the probability density of values occurring in a data set. The probabilities are retrieved by the *ksdensity* function in matlab. All violins are scaled identically making all probabilities comparable.

#### 2. Supplementary Figures

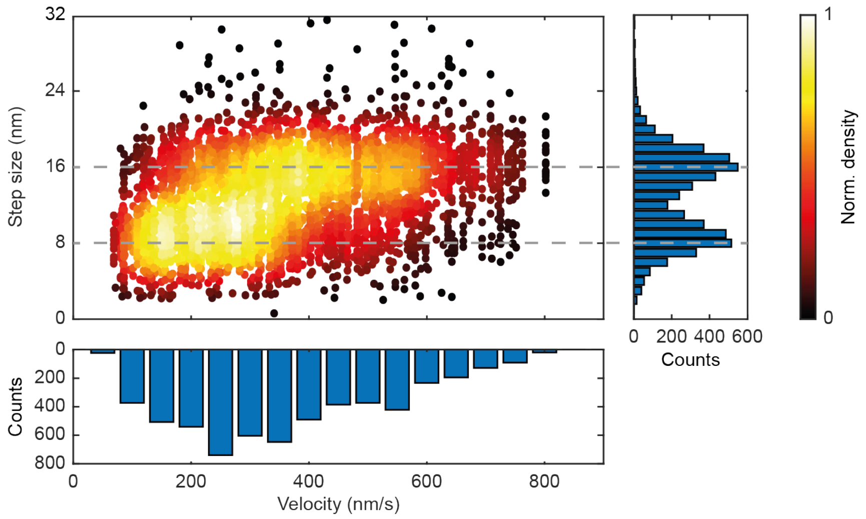

**Fig. S1 Step sizes vary with motor velocity.** (Top left) Bivariate density scatter plot of recorded step sizes over the average velocity of the corresponding trace. The color coding denotes the number of points within 4 nm step sizes and 50 nm/s velocity. Step sizes of 8 nm and 16 nm are highlighted by grey dashed lines. (Top right) Histogram of the step sizes shown in the scatter plot. (Bottom) Histogram of the velocities shown in the scatter plot. The y axis is inversed. All data shown was recorded with construct K28C.

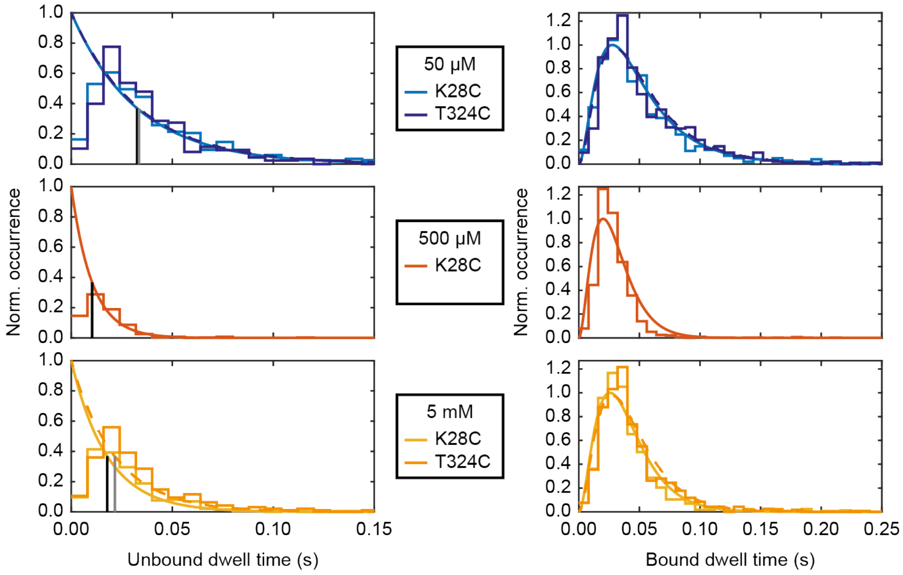

**Fig. S2 Histogram of dwell times in the bound and unbound state.** Stair plots of the dwell time histogram counts in the unbound state (left) and bound state (right) together with the fits according to a single exponential decay (unbound) and a convolution of three exponential decays with two rate constants. The data is split by the employed ATP concentration (top 50 µM, middle 500 µM, bottom 5 mM) and constructs (K28C light blue and yellow, T324C dark blue and orange). For visibility, the fits for construct T324C are shown as dashed lines.

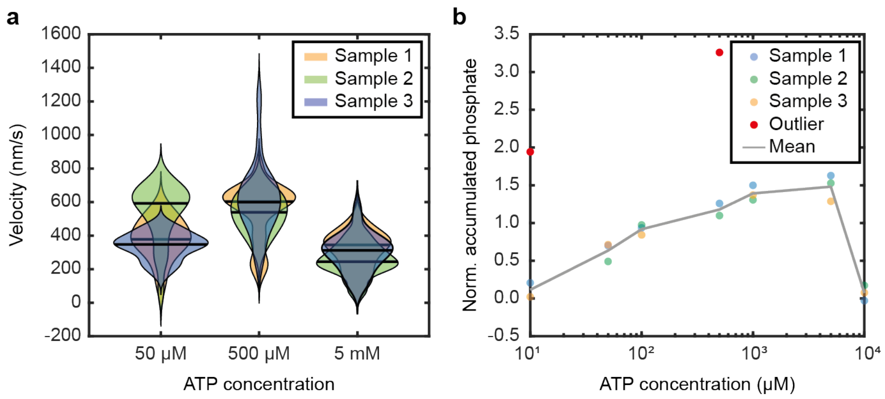

**Fig. S3 Activity controls for construct K28C.** **(a)** Violin plots of the motor velocity measured at 50 µM, 500 µM and 5 mM ATP concentration using a widefield fluorescence microscope. The width of the violin denotes the probability of a certain velocity and the black line the median. For each sample and ATP concentration, the velocity of 20 traces was recorded. **(b)** ATP phosphate assay measuring the amount of phosphate produced by kinesin motors at various ATP concentrations. The amount of phosphate produced at 0 µM ATP was subtracted as a baseline. For each concentration three replicates/ samples were measured. The red circles denote outliers caused by dirt in the sample or crystallization in the buffer medium. The mean values (after removing the outliers) are connected by a grey line.

#### 3. Supplementary Tables

**Table S1. MINFLUX tracking parameters.** $L$-values, laser power (as measured going into the galvo scanner) and repeats used in the different MINFLUX steps

| Step | $L$ (nm) | Laser power (μW) | repeats |
|---|---|---|---|
| 1 | 240 | 6 | 3 |
| 2 | 120 | 24 | 5 |
| 3 | 70 | 70 | 10 |
| 4 | 50 | 180 | 1482 |

### 4. References

1. C. M. Gurth *et al.*, Neurofilament Levels in Dendritic Spines Associate with Synaptic Status. *Cells* **12**,  (2023).
2. A. Yildiz, M. Tomishige, R. D. Vale, P. R. Selvin, Kinesin Walks Hand-Over-Hand. *Science* **303**, 676-678 (2004).
3. M. Tomishige, R. D. Vale, Controlling kinesin by reversible disulfide cross-linking. Identifying the motility-producing conformational change. *J Cell Biol* **151**, 1081-1092 (2000).
4. J. Schindelin *et al.*, Fiji: an open-source platform for biological-image analysis. *Nat Methods* **9**, 676-682 (2012).
5. J. O. Wolff *et al.*, MINFLUX dissects the unimpeded walking of kinesin-1. *Science* **379**, 1004-1010 (2023).
6. R. Killick, P. Fearnhead, I. A. Eckley, Optimal Detection of Changepoints With a Linear Computational Cost. *J Am Stat Assoc* **107**, 1590-1598 (2012).
